# Accurate Protein Function Prediction via Graph Attention Networks with Predicted Structure Information

**DOI:** 10.1101/2021.06.16.448727

**Authors:** Boqiao Lai, Jinbo Xu

**Affiliations:** Toyota Technological Insititute at Chicago, Chicago, IL 60637, USA

## Abstract

Experimental protein function annotation does not scale with the fast-growing sequence databases. Only a tiny fraction (<0.1%) of protein sequences in UniProtKB has experimentally determined functional annotations. Computational methods may predict protein function in a high-throughput way, but its accuracy is not very satisfactory. Based upon recent breakthroughs in protein structure prediction and protein language models, we develop GAT-GO, a graph attention network (GAT) method that may substantially improve protein function prediction by leveraging predicted inter-residue contact graphs and protein sequence embedding.

Our experimental results show that GAT-GO greatly outperforms the latest sequence- and structure-based deep learning methods. On the PDB-mmseqs testset where the train and test proteins share <15% sequence identity, GAT-GO yields Fmax(maximum F-score) 0.508, 0.416, 0.501, and AUPRC(area under the precision-recall curve) 0.427, 0.253, 0.411 for the MFO, BPO, CCO ontology domains, respectively, much better than homology-based method BLAST (Fmax 0.117,0.121,0.207 and AUPRC 0.120, 0.120, 0.163). On the PDB-cdhit testset where the training and test proteins share higher sequence identity, GAT-GO obtains Fmax 0.637, 0.501, 0.542 for the MFO, BPO, CCO ontology domains, respectively, and AUPRC 0.662, 0.384, 0.481, significantly exceeding the just-published graph convolution method DeepFRI, which has Fmax 0.542, 0.425, 0.424 and AUPRC 0.313, 0.159, 0.193.

## 1. Introduction

High throughput sequencing technology has yielded an explosive number of sequences, but only a tiny fraction of them have experimentally determined functional annotations[1]. There is a dire need for efficient and accurate protein function annotation tools for the community to study the growing sequence databases[2–4]. Many computational methods have been developed to annotate protein functions based on primary sequences[5–9], protein family and domain annotations[10–12], protein-protein interaction(PPI) networks[7, 9, 10], and other hand-crafted features[8, 9, 13]. Critical Assessment of Functional Annotation(CAFA), a community-driven benchmark effort for automated protein function annotation, has shown that integrative prediction methods that combine multiple information sources usually outperform sequence-based methods[2–4]. Sequence-based methods use sequence similarity to transfer functional information and thus, do not work well on novel sequences that are not similar to any annotated sequences[2, 4, 14]. Domain and family annotations as well as PPI (Protein-Protein Interaction) information are useful for function prediction, but they are often missing or incomplete for the vast majority of unannotated sequences[12, 15].

Proteins acquire their function by folding into certain 3-dimensional structures *in vivo* [16, 17]. Two structurally similar proteins may share similar functions even with dissimilar sequences [18–21]. That is, purely sequence-based approaches may not work well in transferring functions between structural homologues. To bridge the gap between sequence and function, it is crucial to develop methods that can directly utilize structural information for function prediction. Methods that leverage protein structure databases such as Funfam and DeepFRI[15, 22] have shown promising results in structure-based protein function annotation. While only a very small percentage of proteins have experimental structures, recent breakthroughs in protein contact and structure prediction[23–25] allow us to generate accurate structure information for a large portion of proteins, which can be used for large-scale automated protein function annotations.

Deep learning such as convolutional neural networks(CNN)[26] and residual neural networks(ResNet)[27] has been widely adopted by the computational biology community and showed immense success in some areas such as profiling epigenomic landscapes from DNA sequences[28–31] and protein folding[23, 24, 32]. Graph convolutional networks(GCNs)[33] are able to learn representation from arbitrarily structured graph input[34, 35]. Graph attention network(GAT)[36] is a type of graph neural network (GNN) that performs graph convolution with self-attention[37]. GAT and GNN are used to model gene expression and study protein structure refinement[38, 39]. Unsupervised protein sequence models are used to capture inter-residue relationships for protein contact prediction and have become an integral part of many protein structure prediction methods[23, 24, 32]. Recently, deep protein language models have been developed[40–42] to encode context and global information of a protein for downstream tasks such as stability prediction.

To leverage predicted structure information and protein embeddings for function prediction, we have developed a GAT-based method called GAT-GO. We use RaptorX[43] to predict structure information of a protein sequence and Facebook’s ESM-1b[40] to generate residue-level and sequence-level embeddings of a protein sequence. GAT-GO outperforms traditional homology-based algorithms such as BLAST[44] and previous deep learning methods [6], even when test proteins have low sequence identity with training proteins. Two recent studies [15, 45] have explored GCN and protein embeddings for protein function prediction, but they show limited improvement over sequence-only methods. Our method differs from the GCN method DeepFRI [15] as follows. We use GAT [36] instead of conventional GCN. GAT enhances model capacity by allowing flexible node feature aggregation through a self-attention mechanism. In addition, we use topological pooling[46] to enable more efficient downsampling that improves model generalizability. Moreover, GAT-GO uses predicted inter-residue contacts for both training and test, while DeepFRI uses some native contact graphs in training.

## 2. Results

### 2.1 GAT-GO: predicting protein function via graph attention networks

As shown in Fig. 1, GAT-GO integrates protein sequence representations and predicted inter-residue contact graphs using a CNN-based feature encoder and a GAT-based graph encoder. GAT allows capturing interactions among spatially close residues, which may be missed in sequence-only methods. GAT-GO consists of three major modules: 1) a convolutional neural network(CNN) that takes sequential features and residue-level sequence embedding as input to produce per residue feature representation. 2) a GAT that takes a predicted contact graph and the CNN-generated representation vector as input. Each GAT layer is followed by an attention-based topological pooling layer[46] to perform topology-aware downsampling. To extract protein-level representation, a global pooling layer is used at the end of GAT. 3) a dense classifier that predicts the probability of functional annotations from the representation generated by GAT jointly with protein-level sequence embedding. By default, GAT-GO uses RaptorX-predicted contact graphs and protein embeddings generated by ESM-1b.

**Fig.1.**
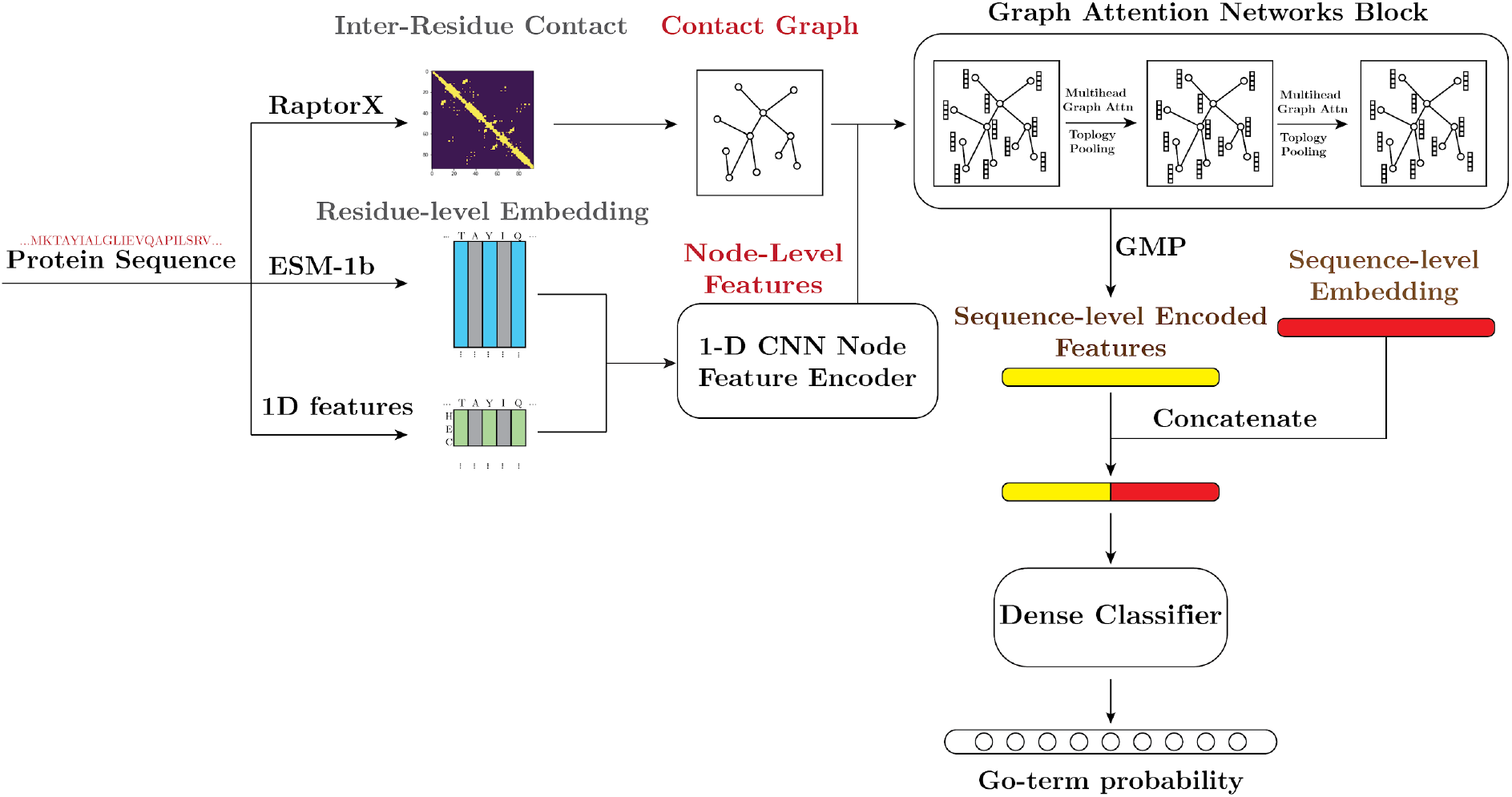
Overall architecture of GAT-GO: Input sequences are first processed into 1-dimensional features(SA/SS/PSSM) and fed into a 1-dimensional CNN feature encoder to produce node-level feature embeddings. Graph Attention Networks(GAT) combines the inter-residue contact graphs with node-level feature embeddings to generate a sequence-level feature vector, which is then used by a dense classifier to predict GO term probability.

### 2.2 GAT-GO improves protein function prediction

We evaluate GAT-GO on the PDB-cdhit dataset and compare it to sequence-only methods including BLAST and the standard Naive baseline used in the CAFA benchmark[2, 4] and a just-published structure-based GCN method DeepFRI. Here DeepFRI uses contact graphs extracted from native structures. We have also implemented a 1D CNN method to represent the state-of-the-art sequence-only deep learning method DeepGO[6]. We evaluate the performance across all three gene ontology domains (MFO, BPO, CCO) using both protein-centric metric*F*_*max*_ (Maximum F score) and GO-term-centric metric *AUPRC*(Area Under Precision-Recall Curve). AUPRC measures the precision-recall tradeoff in a label imbalanced environment. As shown in Table 1, GAT-GO vastly outperforms BLAST, 1D CNN, and DeepFRI across all three gene ontology domains. GAT-GO has *F*_*max*_ 0.637, 0.510, 0.542 on the MFO, BPO, CCO ontology domains, respectively, while BLAST has *F*_*max*_ 0.497, 0.399, 0.390. Despite using predicted contacts and being trained on a much small dataset, GAT-GO has AUPRC 0.662, 0.384, 0.481 on the MFO, BPO, CCO ontology domains, respectively, substantially better than DeepFRI that has AUPRC 0.313, 0.159, 0.193. Here, we evaluate the methods in terms of both the protein-centric *F*_*max*_ metric that measures how methods retrieve relevant function annotations across all tested proteins and the term-centric *AUPRC*metric that assesses the trade-off between the true positive rate and positive predictive value across all GO terms predicted. That is, the higher the *AUPRC*value, the more confidence we have in the predicted protein-function pairs. We observed that GAT-GO has a much larger advantage over the competing methods in terms of *AUPRC*, which indicates GAT-GO predicted function annotations are much more reliable than other methods across all predicted GO terms.

**Table 1:** *F*_*max*_ and *AUPRC*of competing methods on the PDB-cdhit dataset. DeepFRI(Native) indicates the native contact map of a test protein is used.

| Model | $F_{max}$ | | | AUPRC | | |
| --- | --- | --- | --- | --- | --- | --- |
|  | MFO | BPO | CCO | MFO | BPO | CCO |
| Naive | 0.156 | 0.244 | 0.318 | 0.075 | 0.131 | 0.158 |
| BLAST | 0.498 | 0.400 | 0.398 | 0.120 | 0.120 | 0.163 |
| 1D CNN(DeepGO) | 0.359 | 0.295 | 0.420 | 0.368 | 0.210 | 0.302 |
| DeepFRI(Native) | 0.542 | 0.425 | 0.424 | 0.313 | 0.159 | 0.193 |
| GAT-GO(Ours) | **0.633** | **0.492** | **0.547** | **0.660** | **0.381** | **0.479** |

### 2.3 GAT-GO’s performance with respect to sequence identity

Past studies often use time-gated temporal datasets to evaluate model generalizability to previously unannotated sequences [6–8]. However, this temporal evaluation approach may have similar sequences in the training and test data and thus, inflate the test result. Some recent studies use sequence identity to split the training and test data [15, 45] and thus, may provide a more accurate view of model generalizability to novel sequences. This evaluation approach is widely adopted by other fields such as protein structure prediction[47]. We generate training and test datasets (See Methods 4.1) using five different sequence identity thresholds(15%, 25%, 35%, 45%, 55%) and compare GAT-GO with BLAST and two sequence-based methods implemented by ourselves: 1) a 1D CNN method that predicts protein function from primary sequence only; 2) a 1D ResNet method that predicts protein function from both sequential features and protein sequence embeddings. Neither 1D CNN nor 1D ResNet uses predicted inter-residue contact graphs. Here we could not compare GAT-GO with DeepFRI since the latter does not have pre-trained models for a specific sequence identity threshold. As shown in Fig. 2, the performance of almost all the test methods increases with respect to sequence identity, but BLAST performs badly at low sequence identity zones. GAT-GO consistently outperforms the other methods on all three ontology domains regardless of sequence identity. When the training and test sequences share ≤15% sequence identity, GAT-GO has*F* _*max*_ 0.501, 0.406, 0.508 for the MFO, BPO, CCO ontology domains, respectively, and AUPRC 0.427, 0.253, 0.411, much better than 1D ResNet (*F*_*max*_ 0.408, 0.331,0.450 and *AUPRC*0.294, 0.184, 0.318) and 1D CNN (*F*_*max*_ 0.154, 0.161, 0.270 and AUPRC 0.048, 0.067, 0.112). The sequence-only methods 1D CNN and BLAST do not fare well at low sequence identity due to a lack of explicitly shared sequence patterns between the training and test sequences. On the other hand, structural features like predicted inter-residue contact graphs and protein sequence embedding can drastically improve protein function prediction for novel sequences.

**Fig.2:**
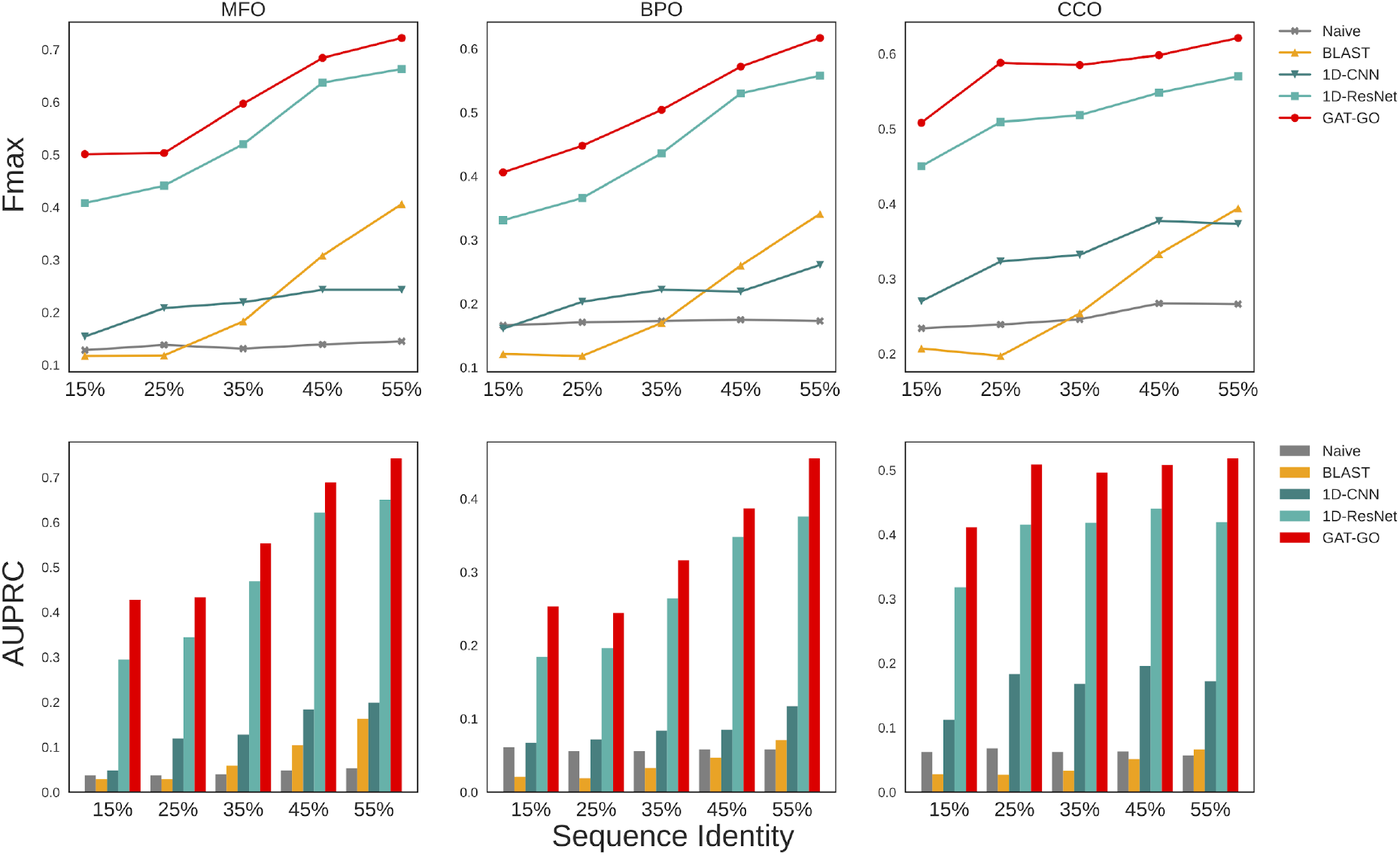
(A)Fmax and (B)AUPRC performance comparison on the PDB-mmseq dataset across different sequence identity thresholds

### 2.4 Predicted contacts and sequence embedding improve protein function prediction

To thoroughly investigate the contributions of individual factors, we evaluate the performance of four deep models (1D CNN, 1D ResNet, GCN, and GAT) with different feature combinations on the PDB-cdhit testset. As shown in Table 2, both 1D ResNet and GAT-GO can leverage protein sequence embeddings to improve function prediction. For example, compared to 1D ResNet using only primary sequence, 1D ResNet using both primary sequence and protein-level embeddings may improve *F*_*max*_ by 0.204, 0.136, 0.109 for the MFO, BPO, CCO ontology domains, respectively. This observation is consistent with[45] that found protein-level sequence embeddings encode useful information for protein function prediction and can significantly enhance model performance.

**Table 2.** *F*_*max*_ and *AUPRC*of different methods with different feature combinations on the PDB-cdhit dataset.

| Model | Input Features | $F_{max}$ | | | AUPRC | | |
| --- | --- | --- | --- | --- | --- | --- | --- |
|  |  | MFO | BPO | CCO | MFO | BPO | CCO |
| 1D CNN | Primary sequence | 0.344 | 0.268 | 0.423 | 0.355 | 0.197 | 0.290 |
| 1D ResNet | Primary sequence | 0.337 | 0.284 | 0.369 | 0.334 | 0.188 | 0.267 |
|  | Primary sequence & protein-level embeddings | 0.541 | 0.416 | 0.478 | 0.541 | 0.289 | 0.386 |
|  | Sequential features & pre-trained embeddings<sup>1</sup> | 0.548 | 0.416 | 0.500 | 0.559 | 0.293 | 0.393 |
| GCN | Predicted contact & residue-level embeddings | 0.459 | 0.443 | 0.461 | 0.408 | 0.245 | 0.315 |
| GAT-GO | Predicted contact & residue-level embeddings | 0.551 | 0.472 | 0.490 | 0.558 | 0.289 | 0.364 |
|  | Predicted contact & sequential features & residue-level embeddings | 0.567 | 0.481 | 0.495 | 0.565 | 0.301 | 0.380 |
|  | Predicted contact & sequential features & pre-trained embeddings | **0.637** | **0.501** | **0.542** | **0.662** | **0.384** | **0.481** |

1. Pre-trained embeddings refer to both residue-level and protein-level pre-trained embeddings (See Methods 4.2)

To investigate the contribution of protein-level embeddings to the performance of GAT-GO, we compare two GAT models. One uses sequential features and residue-level embeddings as input and the other uses both protein-level and residue-level embeddings on top of sequential features. Using protein-level embeddings improves *F*_*max*_ by 0.070, 0.020, 0.052, and *AUPRC*by 0.097, 0.083, 0.101 for the MFO, BPO, CCO ontology domains, respectively. To demonstrate the improvement of the GAT-based model architecture over the GCN-based architecture developed by [15], we compare a GAT model that uses one-hot encoded primary sequences, residue-level embeddings, and predicted inter-residue contacts with a GCN model that uses the same set of features. The GAT model yields *F*_*max*_ of 0.551, 0.472, 0.490, and *AUPRC*of 0.558, 0.289, 0.364 while the GCN model yields *F*_*max*_ of 0.459, 0.443, 0.461, and *AUPRC*of 0.408, 0.245, 0.315 for the MFO, BPO, CCO ontology domains, respectively.

To study how much predicted inter-residue contacts may help, we compare two models. One is a ResNet model that uses sequential features and both protein-level and residue-level embeddings but not predicted contact graphs. The other is a GAT model that uses sequential features and both protein-level and residue-level embedding in addition to predicted contact graphs. The GAT-based structure-aware model has *F*_*max*_ 0.637, 0.501, 0.542, and AUPRC 0.662, 0.384, 0.481 for the MFO, BPO, CCO ontology domains, respectively, while the ResNet model has*F*_*max*_ 0.548, 0.416, 0.500 and AUPRC 0.559, 0.293, 0.393. See Supplementary Table S2-S3 for more detailed comparisons. This result suggests that predicted structural information can improve protein function prediction in addition to protein sequence embeddings.

### 2.5 Explicit Structural Information Amends Function Interpretation in Longer Sequences and High-Specificity GO Terms

To better understand how predicted inter-residue contacts enhance protein function prediction, we compare GAT-GO with ResNet, both using the same set of input features except that ResNet does not use predicted contact graphs. To measure how GAT-GO improves function prediction accuracy, we calculate the precision difference of GAT-GO and ResNet on each tested sequence and study its relationship with sequence length. The precision improvement by GAT-GO is positively correlated with sequence length. Their spearman’s correlation coefficient is 0.312, 0.259, 0.221 for the MFO, BPO, CCO ontology domains, respectively, as shown in Fig. 3. This result implies that the longer the sequence, the greater impact predicted structural information has on the prediction accuracy. This is because GAT (with predicted contacts) may capture interactions among residues that are well separated along the primary sequence. In contrast, sequence-based methods such as 1D ResNet often focus on local sequence patterns and are not good at capturing interactions of residues that are far away from each other along the primary sequence. Studies have shown that explicitly modeling long-range residue interactions can greatly improve performance in various tasks[38] in addition to functional prediction.

**Fig.3:**
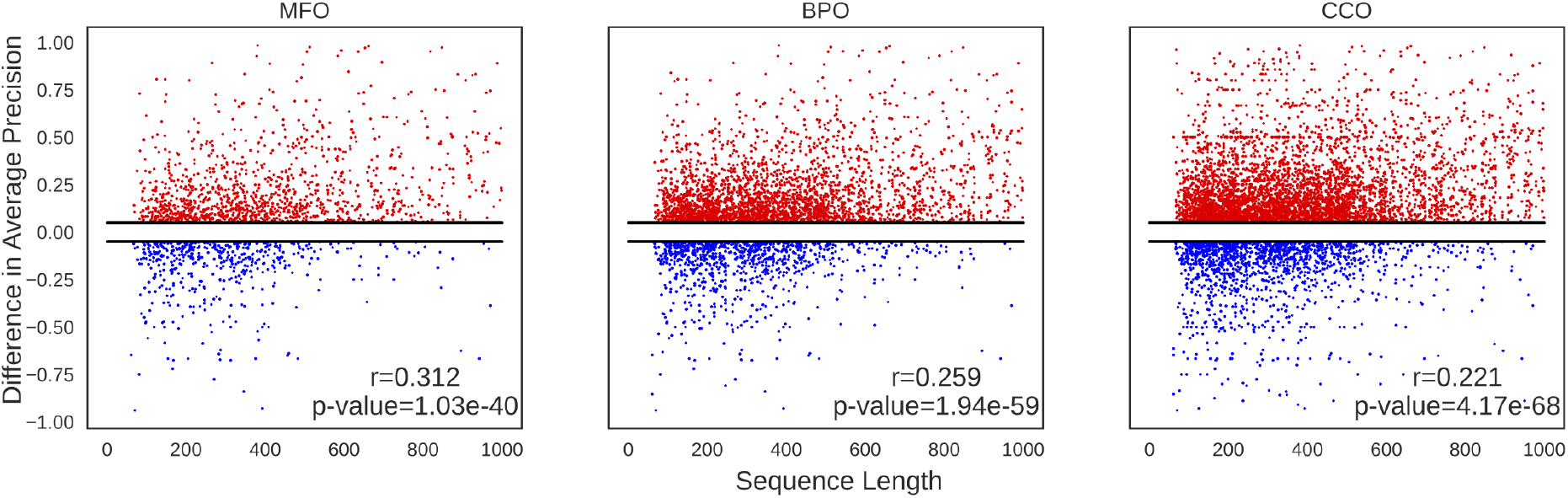
Performance improvement of GAT-GO over the ResNet-based sequence method vs. sequence length. Sequences with increased average precision (red) and decreased average precision(blue) are plotted against the length of the sequences. Spearman’s correlation coefficients for the MFO, BPO, CCO ontology domains are 0.312, 0.259, 0.221 with p-value of 1.03e-40, 1.94e-50, 4.17e-68 respectively. Where the two-sided p-values are calculated under the student’s t distribution with n-2 degree of freedom.

Using predicted structural information may also improve the prediction accuracy of GO terms with high specificity. We measure the specificity of one GO term using information content (See methods 4.5). A Go term with high information content (IC) is more specialized and thus, rarer in occurrence. On the other hand, a GO term of low information content represents a broad function that is more common in annotations. As shown in Fig.4, GO terms with IC>12 benefit more from the inclusion of inter-residue contact graphs than those with IC<6.

**Fig.4:**
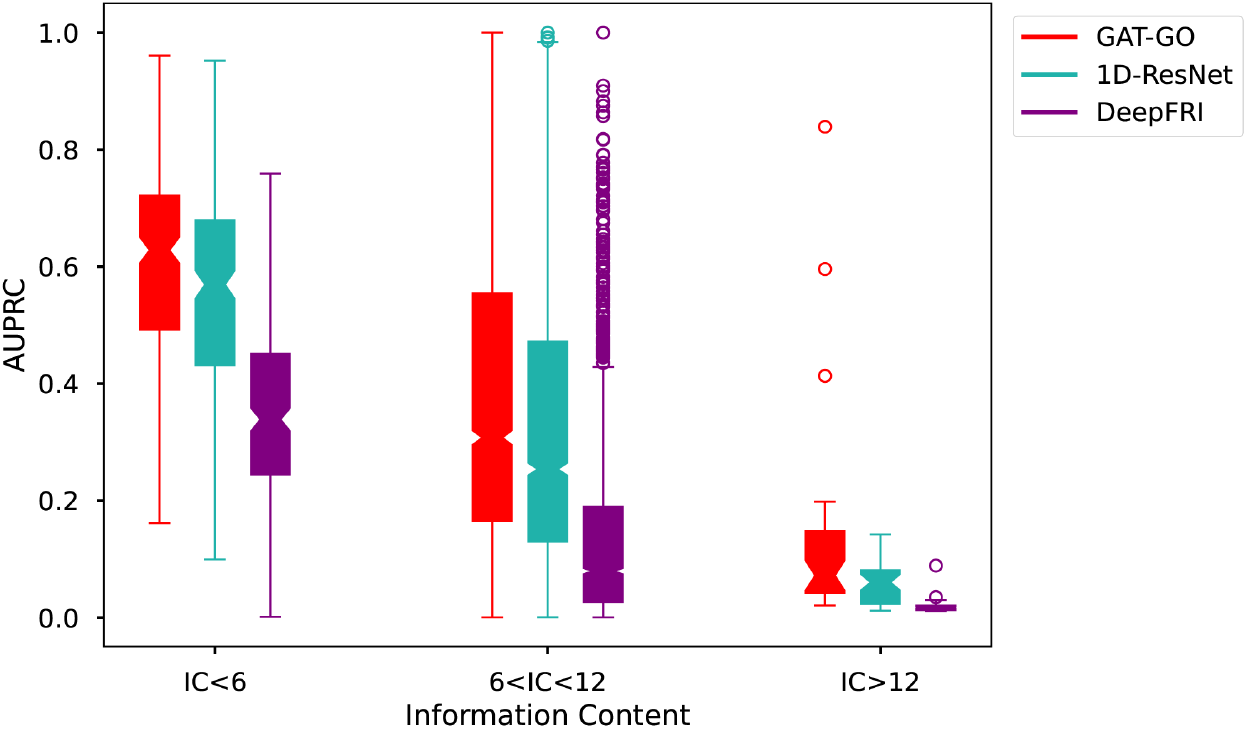
Distribution of AUPRC score on GO-Terms covered by GAT-GO over different specificity levels (Information Content) for different methods: GAT-GO(red), 1D-ResNet(green), and DeepFRI(purple). The box representation has the center as the median, upper and lower edges are the interquartile range and the whiskers are the data range.

## 3. Discussion

In this study, we have presented GAT-GO, a structure-based deep learning method that integrates predicted inter-residue contact graphs, protein embedding, and sequential features for protein function prediction. By integrating predicted contacts and protein embedding through graph attention networks, our method may accurately and efficiently map protein sequences to function annotation at a large scale, especially when the test sequences are not similar to the training sequences. The combination of sequential features, protein embeddings, and inter-residue contact graph leads to one of the major advantages of GAT-GO which is its capability to jointly predict protein function from both local and global information. In contrast, sequence-based methods cannot make use of predicted structure information and thus, are not good at handling a test sequence that is not similar to any training sequences.

GAT-GO outperforms existing function prediction methods including the recently developed structure-based method by utilizing high-resolution structure information and high-capacity pre-trained protein embeddings. Our experiment shows that both protein embeddings and predicted contact graphs, as well as the GAT-based model architecture, can significantly improve function prediction. To further improve structure-based protein function annotation, instead of using predicted contact graphs, we may use predicted inter-residue distance graphs or 3D structure coordinates as the structure representation. Other network-based protein features such as protein-protein interaction (PPI) networks may also be used.

## 4. Methods

### 4.1 Data Split and Processing

We download the data used by DeepFRI[15] at https://github.com/flatironinstitute/DeepFRI. We denote it as PDB-cdhit since the train/test split is generated by cd-hit[48] with 40% sequence identity. Each sequence is annotated with memberships of 2,752 GO terms across three ontology domains. CD-hit[48] is often used to remove redundancy between training and test proteins, but its greedy clustering method and local-alignment-based sequence similarity calculation can still lead to redundancy between the training and test set and thus, inflate the test results[48]. To fix this, we employ another sequence clustering tool MMseqs to generate a new dataset denoted as PDB-mmseqs. MMSeqs uses a non-greedy clustering scheme and profile-based alignment method to ensure there is no higher than desired sequence identity across clusters[49]. To split data by sequence identity, we use MMseqs to cluster all protein sequences with a given sequence identity threshold, and then select a representative sequence from each cluster to build a seed sequence pool. Then we split the seed sequence pool uniformly at random with an 8/2 ratio to form the train and test seed sets. The final train/test sequences are determined by including sequences from the respective clusters of the train/test seeds. Validation sequences are generated by sampling 10% of the sequences from the training set uniformly at random. We generate 5 different data splits with 5 different sequence identity thresholds (15%, 25%, 35%, 45%, 55%). That is, in each dataset, no training and testing protein share higher than the respective sequence identity. See Supplementary Table S1 for details of the datasets.

To measure the information content for an individual Go term, we compute the Shannon Information from its frequency in the training set. See Supplementary Section 5 for more details.

### 4.2 Input Features

#### Sequential features

One-hot encoding of the primary protein sequence is the most commonly used input feature for sequence-based methods. We encode a sequence with 25 different symbols including the 20 common amino acid symbols and 5 compound ambiguous symbols. We also use position specific scoring matrix (PSSM) as sequence profiles derived from the profile HMM generated by HHblits[50] with E-value=0.001 and uniclust30 dated in August 2018. We use secondary structure and solvent accessibility predicted by RaptorX-Property[51] from PSSM. In the article, sequential features are referred to as the combination of one-hot encoded primary sequence, sequence profile, secondary structure annotations, and solvent accessibility annotations.

#### Predicted protein structure information

To obtain inter-residue contact graphs, we predict protein Cb-Cb distance using RaptorX[23] and define inter-residue contact probability as the probability of the Cb-Cb distance < 8 Å. To build the contact graph, we add an edge between two residues if and only if they have >50% predicted contact probability. We have evaluated performance with respect to different contact probability cutoffs in Supplementary Table S4.

#### Protein embeddings

To obtain residue-level sequence embeddings, we use ESM-1b[40], a transformer-based self-supervised protein language model trained on the UniRef UR50/50 database[52]. We generate the protein-level embedding from the residue-level embedding by averaging across all residue positions.

### 4.3 Graph Attention Networks

Graph Neural Network (GNN) is a powerful tool for extracting information from arbitrarily structured graph data[53]. Graph Convolutional Network (GCN) uses spectral convolution on the graph Fourier domain to aggregate neighboring representation for feature learning[33]. In this study, we use the first-order approximation of the spectral convolution: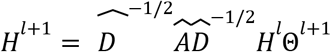

Where 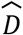 is the degree matrix such that 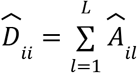, and 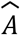 is the adjacency matrix of the input contact graph with self-loops, i.e., 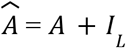 where *I*_*L*_ is an *L* × *L* identity matrix. *H*^*l*^ is the graph representation at layer *l*, and Θis the trainable weight of the neural network.

Graph Attention Networks (GAT) aim to improve the flexibility and capacity of the graph spectral convolution by employing a self-attention mechanism to parameterize the feature aggregation process[36]. Compared to GCN which aggregates neighboring features with fixed weights determined by the degrees of the respective nodes, GAT uses learnable self-attention-based weights. First, GAT calculates the pairwise importance scores for all node pairs as*e*_*ij*_ = *a*(Θℎ_*i*_, Θℎ_*j*_)where ℎ^*l*^_*i*_ is the hidden representation for node *i* at layer *l*and Θ^*l*^ is the trainable weight for layer *l*. We use a single-layer feedforward neural network as our attention function, i.e., *a*(*Z*_*i*_, *Z*_*j*_) = *LeakyReLU*(*w*^*T*^[*Z*_*i*_ ||*Z*_*j*_]) where *w* is the trainable attention weight vector and ||denotes concatenation. Graph structure information is then injected by using the mask attention score α_*ij*_ ∀ *j* ∈ *N*_*i*_ on top of the pairwise importance score with softmax as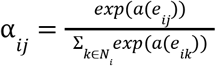, where *N*_*i*_is the neighborhood of node *i*. The hidden representation of each node is then updated with 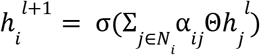 where σ(·) = *ReLU*(·)is the activation function.

### 4.4 GAT-GO network architecture

GAT-GO consists of three major components: 1) a CNN-based sequence feature encoder. It takes three sequence features as input and encodes them into residue-level feature vectors. The three input features are the one-hot encoding of primary sequences, sequence profile, and residue-level sequence embedding derived from the protein language model. A 512-channel CNN is used to encode each input feature and then summed to generate the final encoding. 2) A GAT-based graph encoding module comprised of 4 GAT layers with 512, 512, 1024, 1024 hidden layers and 12 attention heads with a 0.5 dropout rate. Following each GAT layer, there is a topological pooling layer[46] that computes node-level self-attention scores with graph convolution and the top 50% of the nodes are retained as input to the next GAT layer. A global mean pooling layer is then used to pool residue-level feature encoding into sequence-level feature encoding. 3) A dense classifier that predicts protein function from the sequence-level feature and the sequence-level protein embeddings.

### 4.5 Evaluation Metrics

We evaluate models by two main metrics, protein-centric *F*_*max*_ and GO-term-centric AUPRC (area under the precision-recall curve). *F*_*max*_ is the maximum *F*_1_ score across all prediction thresholds in the range of [0,1] with a step size of 0.01. The AUPRC is a summarization of the precision-recall curve by calculating the weighted mean of the precision achieved at each threshold. We use AUPRC to measure the precision-recall tradeoff in a label imbalanced environment. See Supplementary Section 2 for more details.

### 4.6 Competing Methods

#### DeepFRI

It is a recently published GCN-based method that uses autoregressive protein embeddings and contact graphs derived from 3-dimensional structures[15]. We obtain DeepFRI’s predictions following the instruction at https://github.com/flatironinstitute/DeepFRI using experimentally solved structures downloaded from https://www.rcsb.org/. In contrast, our GAT-GO is a GAT-based method with multi-level topological pooling that uses transformer-based residue- and protein-level embeddings. GAT-GO is trained on inter-residue contact graphs predicted by RaptorX[43] and thus, does not rely on experimentally solved structures. For the purpose of comparing the DeepFRI architecture with different feature combinations, we also implemented a 3-layer GCN model as described in [15] that we could train & test on our custom input features.

#### BLAST

Following the protocol used in the CAFA benchmark[4], we obtain the BLAST score for each sequence by first running blastp against the corresponding training database. For each hit, we transfer the corresponding GO terms with the sequence identity as the predicted probability. When multiple hits contain the same GO term, the maximum sequence identity is retained.

#### CNN and ResNet

We have implemented a 1D CNN model with 16 parallel one-layer convolution operations where each convolution has kernel sizes of [8,16,…,128] and 512 filters. This CNN is very similar to the state-of-the-art sequence-only deep learning method DeepGOplus[6]. We have also implemented a ResNet model with 21 convolution layers that predict protein function from both primary amino acid sequences and sequential features. See supplementary for more details.

### 4.7 Model Training

We train our models with binary cross-entropy as the loss metric and the AdamW optimizer[54] with a learning rate of 1e-4 for 30 epochs. We implemented all models in the Pytorch and Pytorch geometric library[55, 56]. A validation set is used to employ an early stopping scheme with patience of 10 epochs. All models are trained with an NVIDIA 2080Ti GPU on a Linux machine.

