## Supplementary file for "Accurate Protein Function Prediction via Graph Attention Networks with Predicted Structure Information"

Boqiao Lai[1], Jinbo Xu[1,*]

[1]Toyota Technological Insititute at Chicago, Chicago, IL 60637, USA

#### 1. Dataset description

Table S1. Numbers of sequences in the datasets

| Datasets | Number of sequences | | |
|---|---|---|---|
| | Training | Testing | Validation |
| PDB-cdhit | 29,902 | 3,416 | 3,323 |
| PDB-mmseqs | | | |
| 15% | 30,445 | 2,888 | 3,308 |
| 25% | 30,551 | 2,667 | 3,408 |
| 35% | 28,839 | 4,375 | 3,244 |
| 45% | 27,553 | 5,272 | 2,988 |
| 55% | 27,085 | 5,791 | 3,072 |

### 2. Evaluation metrics

#### 2.1 $F_{max}$

$$F_{max} = max_t\left[\frac{2\cdot AvgPr(t)\cdot AvgRc(t)}{AvgPr(t) + AvgRc(t)}\right]$$

$$pr_i(t) = \frac{\Sigma_f I(f\in p_i(t) \land f\in T_i)}{\Sigma_f I(f\in p_i(t))}$$

$$rc_i(t) = \frac{\Sigma_f I(f\in p_i(t) \land f\in T_i)}{\Sigma_f I(f\in T_i)}$$

$$AvgPr(t) = \frac{1}{m(t)} \cdot \sum_{i=1}^{m(t)} pr_i(t)$$

$$AvgRc(t) = \frac{1}{n} \cdot \sum_{i=1}^{n} rc_i(t)$$

Where $f$ is a single GO term, $p_i(t)$ is the binary predictions of GO terms at threshold $t$ for sequence $i$. $T_i$ is the set of ground truth GO terms for sequence $i$. $m(t)$ is the number of sequences that we predict at least one positive GO term at threshold $t$. $n$ is the total number of sequences in the test set. $I(\cdot)$ is the indicator function. $AvgPr(t)$ is the average precision at the threshold $t$ across all test sequences with at least one predicted ontology term. $AvgRc(t)$ is the average recall at the threshold $t$ across all predicted sequences.

#### 2.2 AUPRC

$$AUPRC = \sum_{n=1}^{N} (R_n - R_{n-1})P_n$$

Where $R_n$ and $P_n$ are the recall and precision of the $n^{th}$ threshold and N is the total number of thresholds evaluated. We report the micro-average of the AUPRC where the precision and recall values are computed on all predictions across all tested proteins, then we evaluate AUPRC with the above-mentioned formula. In contrast to the macro-average AUPRC which is the average AUPRC of each individual GO-term across all tested proteins, the micro-averaging approach is more preferable where the labels are imbalanced.

### 3. Sequence-based CNN and ResNet models

#### 3.1 1D CNN

```
Input: one-hot encoded primary sequence
seq_feat = []
for kernel_size in [8, 16, 24, 32, 40, 48, 56, 64, 72, 80, 88, 96, 104, 112, 120, 128]:
    out = Conv1d(input_channel = feat_dim, output_channel = 512, kernel_size = kernel_size)(input)
    out = MaxPool1d(kernel_size = 1000 - kernel_size)(out)
    seq_feat.append(out)
feat = concat(seq_feat)
pred = Dense_Classifier(feat)

Optimizer = AdamW(lr = 1e-4)
```

#### 3.2 1D ResNet

Our 1D ResNet model consists of 8 residual blocks as shown in Fig S1. It takes 1D sequential features as input and first encodes the input feature with a convolution layer of 64 output channels. The 8 residual blocks have channel sizes of 128, 128, 256, 256, 256, 256, 512, 512 respectively. In each block, we have 2 convolution layers with ReLU activation and Instance Norm. Finally, we use one max-pooling layer before feeding it into the dense classifier. Between residual blocks, if the channel size is changed, we apply a downsampling convolution with a kernel size of one to the residual connection for channel size matching.

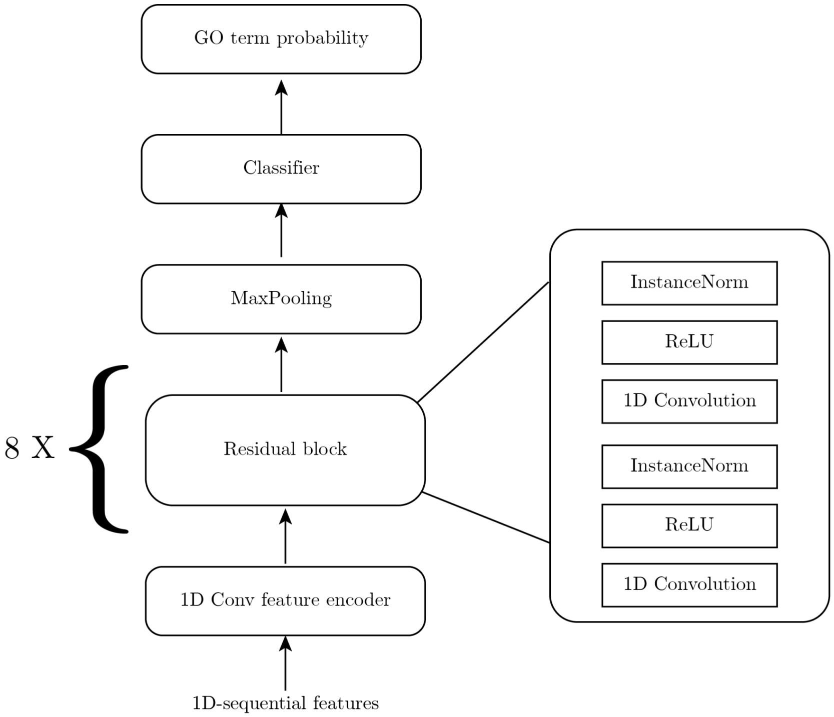


Fig. S1: Overview of the ResNet model.

### 4. Additional performance comparisons

**Table S2.** *$F_{max}$ and AUPRC of sequence-based deep models with different feature combinations on the PDB-chdit dataset.*

| | | $F_{max}$ | | | AUPRC | | |
|---|---|---|---|---|---|---|---|
| Model | Input Features | MFO | BPO | CCO | MFO | BPO | CCO |
| 1D-CNN | Primary sequence | 0.344 | 0.268 | 0.423 | 0.355 | 0.197 | 0.290 |
| | Primary sequence & protein-level embeddings | 0.520 | 0.379 | 0.518 | 0.545 | 0.295 | 0.406 |
| 1D-ResNet | Primary sequence | 0.337 | 0.284 | 0.369 | 0.334 | 0.188 | 0.267 |
| | Primary sequence & protein-level embeddings | 0.541 | 0.416 | 0.478 | 0.541 | 0.289 | 0.386 |
| | sequential features | 0.539 | 0.370 | 0.385 | 0.549 | 0.286 | 0.339 |

| | | | | | | | |
|---|---|---|---|---|---|---|---|
| | sequential features & protein-level embeddings | 0.585 | 0.420 | 0.439 | 0.590 | 0.316 | 0.388 |

**Table S3.** *$F_{max}$ and AUPRC of GAT-GO with different feature combinations on the PDB-cdhit dataset.*

| | | $F_{max}$ | | | AUPRC | | |
|---|---|---|---|---|---|---|---|
| Model | Input Features | MFO | BPO | CCO | MFO | BPO | CCO |
| GAT-GO | Primary Sequence | 0.258 | 0.303 | 0.433 | 0.202 | 0.185 | 0.292 |
| | Primary sequence & residue-level embeddings | 0.551 | 0.472 | 0.490 | 0.558 | 0.289 | 0.364 |
| | sequential features & residue-level embeddings | 0.567 | 0.481 | 0.495 | 0.565 | 0.301 | 0.380 |
| | sequential feature & protein-level embeddings | 0.632 | 0.489 | 0.553 | 0.660 | 0.382 | 0.482 |
| | sequential features & embeddings | 0.637 | 0.501 | 0.542 | 0.662 | 0.384 | 0.481 |

**Table S4.** *$F_{max}$ and AUPRC of GAT-GO under different contact probability cutoffs and batch sizes on the PDB-cdhit dataset.*

| | | $F_{max}$ | | | AUPRC | | |
|---|---|---|---|---|---|---|---|
| Contact Cutoff | Batch Size | MFO | BPO | CCO | MFO | BPO | CCO |
| > 0.3 | 1 | 0.619 | 0.520 | 0.520 | 0.569 | 0.292 | 0.349 |
| | 2 | 0.620 | 0.491 | 0.528 | 0.627 | 0.340 | 0.424 |
| | 4 | 0.615 | 0.487 | 0.537 | 0.626 | 0.342 | 0.450 |
| | 8 | 0.625 | 0.485 | 0.530 | 0.638 | 0.332 | 0.433 |
| > 0.5 | 1 | 0.610 | 0.516 | 0.519 | 0.559 | 0.303 | 0.356 |
| | 2 | 0.627 | 0.514 | 0.538 | 0.618 | 0.326 | 0.399 |
| | 4 | **0.637** | **0.510** | **0.542** | **0.662** | **0.384** | **0.481** |
| | 8 | 0.625 | 0.488 | 0.528 | 0.645 | 0.338 | 0.431 |

### 5. Distribution of IC in the PDB-cdhit training set

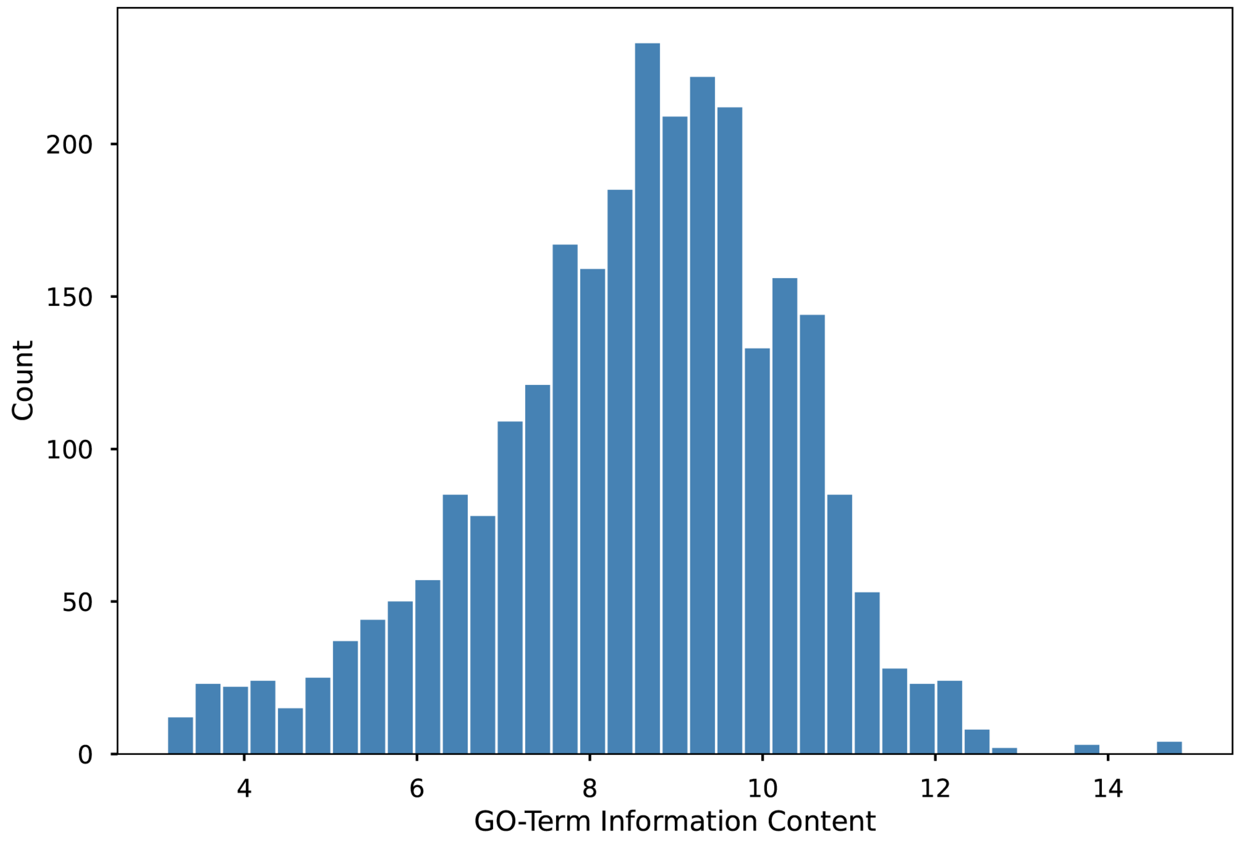


**Fig. S2**. *Distribution of Information Content(IC) for GO-terms in the PDB-cdhit training set. IC is measured in Shannon information. That is, the IC of one GO term i is calculated by* $IC_i = -\log_2(P(GO_i))$ *where* $P(GO_i)$ *is the occurring frequency of this term in the training set.*
